# Space use and resource selection of bobcats in the Appalachian Mountains of western Virginia

**DOI:** 10.1101/831644

**Authors:** David C. McNitt, Robert S. Alonso, Michael J. Cherry, Michael L. Fies, Marcella J. Kelly

## Abstract

Bobcats are an apex predator and a species of socio-cultural importance in the central Appalachian Mountains. Despite their importance, knowledge of bobcat spatial ecology in the region is sparse. We examined space use and resource selection of bobcats in the Appalachian Mountains of western Virginia during 3 biological seasons: breeding (January-March), kitten-rearing (April-September), and dispersal (October-December). We observed sex effects on all space use metrics, with male seasonal areas of use (SAU) approximately 3 times larger than female SAUs and male movement rates 1.5 times higher than females during all seasons. We found no seasonal effect on SAU size for either sex. Female movement rates increased during the kitten-rearing season, and male movement rates increased during the dispersal season. We examined seasonal bobcat resource selection at 2 hierarchical scales, selection of home ranges within the landscape (2^nd^ order) and selection of locations within home ranges (3^rd^ order). Female bobcats exhibited 2^nd^ order selection for higher elevations and deciduous forest and avoidance of fields. Males exhibited 2^nd^ order selection for higher elevations and fields. Male 2^nd^ order selection appears to be driven largely by the spatial distribution of females, which is mediated through the valley and ridge topography of the study area. Sample size precluded 3^rd^ order analysis for females, however males exhibited 3^rd^ order selection for higher elevations, fields, and deciduous forest. Resource selection patterns varied seasonally for both sexes, possibly driven by seasonal shifts in prey availability. Our findings highlight the importance of forested ridges to bobcats in the region. Our findings also illustrate the differences in space use between sexes, which future research efforts should consider. Further research should investigate seasonal shifts in bobcat prey selection, which may further explain the seasonal resource selection shifts we observed, and highlight potential implications for prey species.

## Introduction

An understanding of animal spatial ecology is critical for managing and conserving wildlife populations [1,2]. Knowledge regarding space use and habitat requirements of wildlife provides insight into fundamental ecological processes such as population dynamics, behavioral interactions, and foraging behavior [3-5]. This information can be vital for informing wildlife management decisions and understanding a species role within food webs may vary across space.

Bobcat (*Lynx rufus*) populations are increasing throughout much of their range, both through recolonizing areas of previous extirpation and increasing in abundance where they have persisted [6]. These trends are evident in the central Appalachian Mountains [6]. Although bobcats largely persisted, wolves (*Canis* spp.) and cougars (*Puma concolor*) remain extirpated from ecosystems in the Appalachian Mountains, leaving bobcats as an apex predator in the region, along with black bears (*Ursus americanus*) and coyotes (*Canis latrans*) [8]. Within this guild, bobcats are the only obligate carnivore, can occur at population densities approximately twice that of coyotes [9,10], and consume a diversity of taxa [8]. Thus, bobcats hold potential to influence populations of both prey and competitors, and understanding the spatial ecology of bobcats may be important to predicting spatial variation in top-down forces in these systems. In addition to their ecological importance, bobcats have economic and socio-cultural value, as a furbearer and game species throughout most of the central Appalachian Mountains and are widely recognized by the public. Despite their importance, there is a paucity of information on bobcat spatial ecology throughout the central Appalachian Mountains.

Due to the wide variety of ecosystems in which bobcats occur, there is considerable variability in many facets of their spatial ecology, yet there are certain range-wide patterns driven by their biology. Bobcats are solitary, territorial, and exhibit a polygynous reproductive strategy [11]. Thus, males should seek to maintain large home ranges that overlap the home ranges of multiple females, and defend against other males competing for mates [12]. Females raise young independently, and therefore must maintain home ranges that allow access to sufficient resources to support reproduction, while minimizing energy expenditure and exposure to risk from humans or other predators [12]. As a result, male bobcat home ranges are larger than those of females [13]. Male movement rates are often greater than female movement rates [14,15,16,17], but most information on bobcat movement rates comes from studies that employed VHF telemetry, which often had coarse sampling intervals and poor locational accuracy. More recent studies using GPS telemetry have found varied results [18,19,20], and there is currently a scarcity of published bobcat movement rates from GPS studies. As obligate carnivores with a stalk and ambush hunting mode, [11], prey availability is a key driver of bobcat space use and resource selection. Regional variation in bobcat home range size is likely driven primarily by prey availability; specifically, as prey availability increases, individuals are able to acquire necessary resources in smaller areas [13,21,22]. Bobcats select for areas with higher prey abundance and activity [23]. As ambush predators, concealment cover is a crucial component of prey availability, and bobcats are known to select areas of dense vegetation [21,24,25,26].

Bobcat space use may vary seasonally, which is largely attributed to reproductive behavior and varying prey abundance in regions with pronounced seasonality, thus seasonal space use changes are more common in northern latitudes [15,22,26,27,28,29]. When home range sizes vary seasonally, they are typically smaller during summer months when prey is more available and females remain closer to den sites, and larger during winter months when prey may be less available and males are seeking to maximize breeding opportunities [26,27,29,30]. Although females may have smaller home ranges during the kitten-rearing season, their activity and movement rates often increase during this time, indicating more intensive use of home ranges, most likely associated with providing food to offspring [15,19,31]. Both sexes may increase movement rates during winter months, which has been attributed to breeding behavior and decreased prey availability [15,30]. Habitat selection patterns often shift seasonally as a result of changes in availability of prey and cover [15,24,29].

Understanding bobcat ecology and space use often requires region-specific information. Across regions, bobcats occupy a wide variety of habitat types, often with major differences in prey availability and population abundances of sympatric predators. Region-specific data enable comparisons to distribution-wide patterns, insight into drivers of regional variation, and provide local knowledge to managers. Local understanding of bobcat spatial ecology is essential for effective management of bobcat populations and their impacts on prey.

Information on bobcat spatial ecology is currently sparse in the central Appalachian Mountains. Existing information on bobcat spatial ecology in the region comes from studies conducted during the 1980s in Tennessee and Kentucky, both of which had sporadic monitoring associated with use of VHF telemetry and did not examine habitat selection [32,33]. Existing information on bobcat population and trophic ecology in Virginia comes from noninvasive genetic sampling and examination of stomach contents [9,10,34]. In the eastern portion of their range, most research into bobcat spatial ecology has been conducted in the Southeast and Northeast, yet these areas represent considerably different ecosystems than the central Appalachian Mountains. For example, the central Appalachian Mountains contain the highest elevations in the eastern United States. Thus, there is a need to fill the gap in knowledge regarding bobcat spatial ecology in the central Appalachian Mountains.

We examined home ranges, movements, and resource selection of bobcats in the Appalachian Mountains of western Virginia and investigated the effects of sex and season on these facets of spatial ecology. We predicted that bobcat home ranges in our study area would be comparable to estimates from more northern latitudes, that male home range size and movements would be greater than those of females, and that resource selection patterns would vary between sexes. We also predicted that home range size, movement rates, and resource selection would vary among seasons, due to seasonal variation in both reproductive processes and prey availability.

## Materials and Methods

### Study Area

Our study area encompasses the western half of Bath County, Virginia, adjacent to the border with West Virginia (Figure 1). Bath County is in the Valley and Ridge physiographic province of the Appalachian Mountain range, characterized by parallel, northeast-southwest oriented ridges with narrow valleys interspersed. The repetitive topographical pattern results in largely predictable land cover, with public, forested land on the steep ridges and slopes, and narrow strips of private, low-intensity development and agriculture in the flatter valley bottoms. Bath County is 90% forested land cover, most of which is managed by the US Forest Service and Virginia Department of Game and Inland Fisheries (VDGIF). Elevation ranges from 343 meters to 1363 meters. Average monthly temperature can range from 0.8-25.2 °C, with a mean minimum temperature of 4.7 °C in January and a mean maximum temperature of 31.7 °C in July (National Oceanic and Atmospheric Administration, public data 2012). Average annual precipitation was 97.8 cm (National Oceanic and Atmospheric Administration, public data 2012). The forest structure primarily consists of mature deciduous forest, with common overstory species including oak (*Quercus* spp.), hickory (*Carya* spp.), maple (*Acer* spp.), and tulip poplar (*Liriodendron tulipifera*). Conifers are present in some forest stands, with common species including pine (*Pinus* spp.) and hemlock (*Tsuga* spp.). Other than bobcats, the large carnivore guild includes coyotes and black bears. Common bobcat diet items, based on relative frequency of occurrence, are squirrels (*Sciurus* spp.), voles *(Microtus spp., Myodes gapperi)*, mice (*Peromyscus* spp.), cottontail rabbits *(Sylvilagus* spp.), and white-tailed deer (*Odocoileus virginianus)*[9].

**Figure 1.**
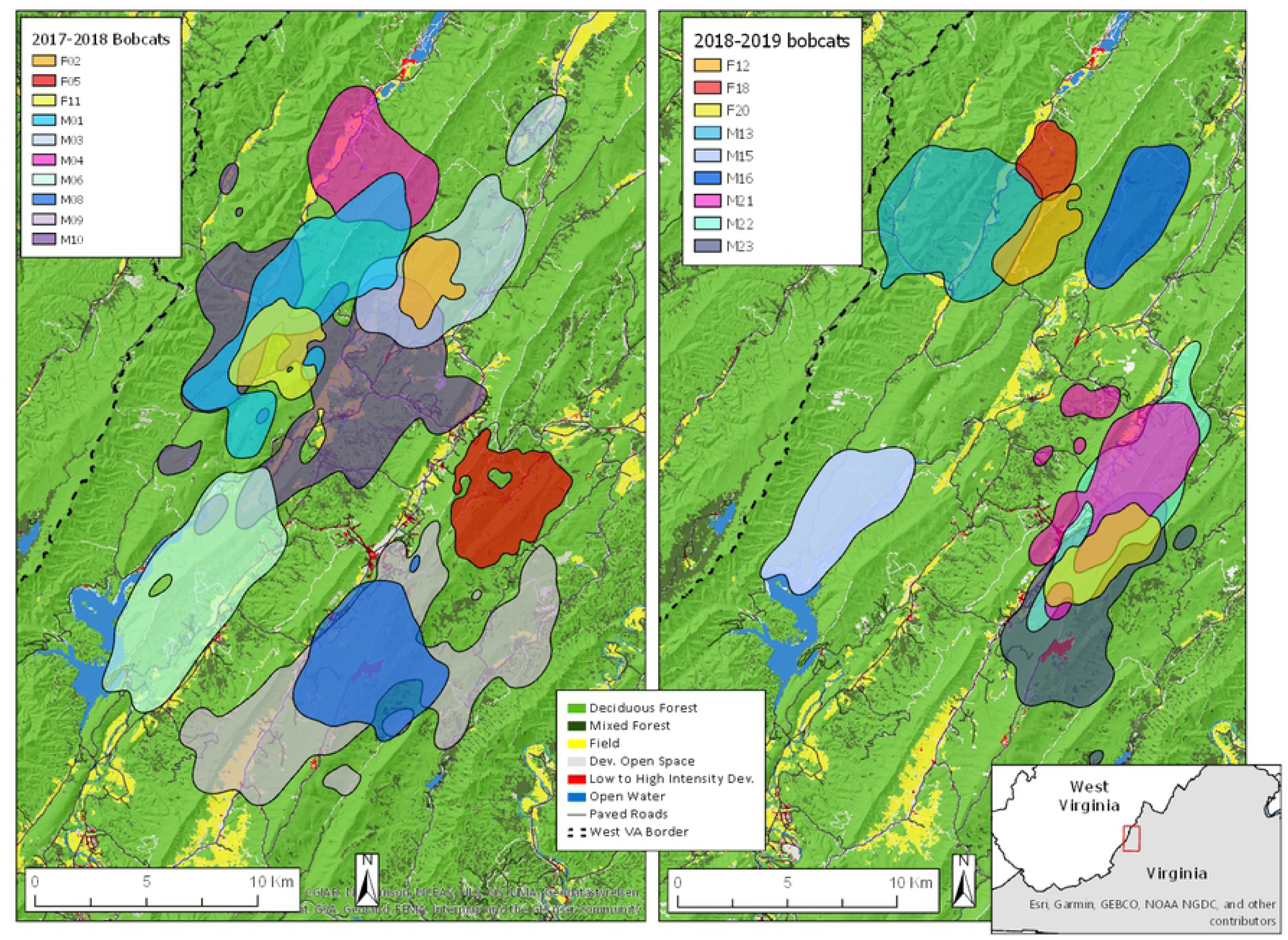
Map of study area with land cover and 95% home ranges of bobcats (n=20) monitored from 2017-2019 in Bath County, VA). Home ranges calculated using the autocorrelated kernel density estimator.

### Bobcat Capture and Monitoring

We captured bobcats using cage traps (Camtrip Cages, Bartsow, California, USA and Briarpatch Cages, Rigby, Idaho, USA) in accordance with Virginia Tech IACUC protocol #16-071. We checked traps twice daily (morning and afternoon). We immobilized bobcats with a mixture of 10-15 mg/kg ketamine hydrochloride and 1mg/kg xylazine using hand injection with syringe. We monitored and recorded respiratory rates, heart rates, and temperatures every 5-10 minutes. We used tooth growth and condition, body morphology, and teat/scrotum characteristics to determine whether bobcats were juvenile or adult [35]. We fitted adult bobcats with Iridium GPS collars (Telonics, Mesa, Arizona, USA and Advanced Telemetry Systems, Isanti, Minnesota, USA). All bobcats captured were marked with color-coded numbered ear tags. Following handling, we reversed xylazine with 0.125 mg/kg yohimbine, administered either rectally or intramuscularly, and allowed bobcats to recover in the cage trap for 30 minutes to 1 hour before release. We programmed GPS collars to collect locations at 1, 2, and 4-hour intervals, however finer-scale sampling was related to other research objectives. For these analyses we used standardized 4-hour intervals.

### Season Classification

We classified 3 seasons of interest based on the reproductive chronology and life history of bobcats. We classified seasons as breeding, kitten-rearing, and dispersal. We classified January 1 – March 31 as the breeding season, to overlap with the estrus cycle of females. We classified kitten-rearing season as April 1 – September 30, since parturition typically peaks in April or May and kittens nurse for up to 4 months [36,37]. We classified the dispersal season as October 1 – December 31. Some previous bobcat space use studies examining seasonal effects have divided the year into only 2 seasons, essentially consisting of breeding (October through April) and kitten-rearing (May through September) periods [14,26,29]. We favor the approach of Chamberlain et al. [15], which divided the year into 3 seasons, breeding, kitten-rearing, and what they termed “winter”, because bobcats are not breeding during autumn and early winter, as evidenced by the lack of parturition during winter. Presumably, resident females will seek to restore body mass depleted during the kitten-rearing period, and resident males will aim to maximize body mass in preparation for the breeding season. Yearling bobcats will likely initiate dispersal during this period. Thus, the months of October through December represent a distinct period of bobcat behavior.

### Home Range Analysis

We estimated bobcat home ranges using the autocorrelated kernel density estimator (AKDE) [38] using the continuous-time movement modeling package (ctmm) [39] in R version 3.5.3 (R Core Team 2019). The AKDE is a third-generation estimator that assumes the data represent a sample from a nonstationary, autocorrelated continuous movement process by incorporating the movement of animals through an autocorrelation function derived from movement models fit to the data [39]. Furthermore, AKDE reduces to a conventional kernel density estimator when locations are truly independent, and can correct for missing locations and irregular sampling schedules through an optimal weighting method [40]. We estimated 95% annual home ranges for bobcats with at least 4 months of relocation data, during at least 2 seasons. We estimated 95% seasonal areas of use (SAU) for bobcats with locations collected for at least 1 month in a given season. We fit linear mixed effects models for each season using restricted maximum likelihood, with area of 95% SAU as the response variables. We used a natural logarithm transformation for home range sizes to meet assumptions of normality. We included both the interaction and main effects of sex and season as predictors, and treated animal-specific intercepts as random effects. We fit models in the package lme4 [42] and assessed the significance of factors and degrees of freedom using Satterthwaite’s method for approximating degrees of freedom in the lmerTest R package [43].

### Movement Analysis

We estimated each bobcat’s movement rates in meters moved per hour, calculated as the straight-line step length between successive locations divided by the time lag. Seasonal movement rates were only examined for bobcats that were monitored for at least one month in a given season. We assigned each step to the appropriate season. We used a generalized linear mixed effects model (GLMM) with a gamma distribution and log link to model movement rates as a function of sex and season. Both the main effects and interaction of sex and season were included as predictors. We treated animal-specific intercepts as random effects. Significance of covariates was determined using Walds Z-test.

### Resource Selection Analysis

We examined seasonal bobcat resource selection at 2 hierarchical scales [5], selection of home ranges within the landscape (2^nd^ order) and selection of locations within home ranges (3^rd^ order), by creating resource selection functions (RSF) in a use-availability framework [1]. We examined 2^nd^ order selection for both sexes, but were only able to examine 3^rd^ order resource selection for males due to the small sample size of females with suitable GPS fix success. Due to poor GPS fix success in some GPS collars, and the potential bias of topography and elevation on GPS fix success, only bobcats with fix success ≥85% were included in 3^rd^ order resource selection analysis.

### Resource Selection Data

For resource selection analysis, we included land cover and topographical based covariates. The land cover covariates we included were distance to deciduous forest, distance to mixed forest, and distance to fields, which we derived from the 30m resolution 2011 National Land Cover Database (NLCD). The deciduous forest covariate consisted of the Deciduous Forest class in the NLCD. To create a mixed forest covariate, we combined the Evergreen Forest and Mixed Forest NLCD classes. To create the field covariate, we combined the Pasture/Hay and Cultivated Crops NLCD classes. Lastly, we created distance raster layers by calculating Euclidian distance to each land cover type using the Euclidean Distance tool in ArcGIS 10.6 (ESRI, Redlands, CA, USA). We used distance-based land cover covariates because they remove the need to base inference on reference categories, reduce the influence of telemetry error, and because effects of land cover types can extend beyond their boundaries (e.g. edge effects surrounding fields). Topographical covariates included elevation and slope at a 30m resolution. We extracted elevation values directly from a digital elevation model (DEM, United States Geological Survey 2013). We calculated slope using the DEM with the Slope tool in ArcGIS 10.6 (ESRI, Redlands, CA, USA).

### 2^nd^ order resource selection

We characterized 2^nd^ order availability following Katnik and Wielgus [44] by simulating random circular polygons. Randomly located, simulated home ranges are superior to landscape proportions for estimating availability. We simulated 10 polygons equivalent in size to each respective bobcat SAU. We then systematically extracted covariate values from every 10^th^ raster cell within both simulated and real SAUs. Instead of simulating polygons throughout the study area for each individual, simulated polygon centers were constrained within a buffer surrounding the centroid of each bobcat’s seasonal locations. We based the buffer size on the space use of 2 dispersing bobcats that we GPS collared. We defined the constraining buffer radius of 5.3 km based on the largest annual home range area of the 2 dispersing bobcats (88.5km^2^), as 5.3 km would be the radius of said home range if it was circular. This area should reflect available habitat more accurately than the entire study area, since it is based on observed area traversed by dispersing bobcats prior to establishing a home range, and habitat on distant portions of the study area may not be realistically available.

### 3rd order resource selection

We examined 3^rd^ order resource selection across seasons only for male bobcats with fix success ≥85% (n=7). To characterize seasonal 3^rd^ order availability, we randomly simulated 10 points within each bobcat’s SAU for every observed GPS location. We constrained availability to SAUs because 3^rd^ order availability may shift seasonally if SAUs change in size or location across seasons. This resulted in 515 ± 107 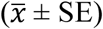 random points per square kilometer. We clipped each bobcat’s observed seasonal locations within SAU borders to remove extraterritorial forays from analysis. We then extracted covariate values at observed locations and simulated points.

### Resource selection model development

To examine 2^nd^ and 3^rd^ order bobcat resource selection, we developed RSFs using binomial generalized linear mixed models (GLMM) in the Program R package lme4 [42]. The binary response variable for 2^nd^ order resource selection was whether a raster cell was extracted from an observed SAU (used = 1) or a simulated polygon (available = 0). The binary response variable for 3^rd^ order resource selection was whether a point was a observed location (used = 1) or a random point (available = 0). Predictor variables for both scales were distance to deciduous forest, distance to mixed forest, distance to fields, elevation, and slope. No covariates were highly correlated (all r < |0.5|; Pearson’s correlation). We included season as an interaction term with all main effects. For 2^nd^ order selection, we created separate models for male and female bobcats, each consisting of the 5 main effects with a season interaction.

For 3^rd^ order selection, we created one model for male bobcats consisting of the 5 main effects with a season interaction. We rescaled all covariates by mean-centering at zero then dividing them by their standard deviation to facilitate model convergence. We included animal-specific random intercepts to account for variation in sampling duration among individuals [45]. We evaluated selection or avoidance based on whether a coefficient significantly differed from zero (α = 0.05). Significance of covariates was determined using Walds Z-test. We inferred selection if used points were closer to habitat variables than random locations, and avoidance if used points were further from habitat variables than random locations.

## RESULTS

We deployed GPS collars on 20 bobcats (14 male, 6 female) from January 2017 through April 2018. Number of locations per bobcat ranged from 259 to 1979, with a mean of 933. Length of collar deployments ranged from 55 – 393 days, with a mean deployment length of 259 days.

We estimated home ranges for 16 resident bobcats (11 males, 5 females) and 2 dispersing males, excluding 2 bobcats (1 male, 1 female) that were monitored for less than 4 months and only during 1 season. We estimated 41 SAUs, including 13 bobcats in the breeding season (8 males, 5 females), 15 bobcats in the kitten-rearing season (11 males, 4 females), and 13 bobcats in the dispersal season (9 males, 4 females). On average, resident male home ranges were 33.9 ± 2.6 km^2^ and were approximately 3 times larger than resident female home ranges (12.1 ± 2.4 km^2^, Figure 2). The home ranges of the 2 dispersing males were 84.8 and 88.5 km^2^ respectively. Male SAUs were larger than female SAUs during all seasons and there was no effect of season on SAU size (Table 1, Figure 2). Average SAU size of females (11.8 ± 1.2 km^2^) and average SAU size of males (32.8 ± 2.0 km^2^) did not differ from average annual home range size of each sex respectively, indicating that SAUs do not shift location extensively throughout the year.

**Table 1.** Linear mixed model for bobcats monitored during 2017-2019 in Bath County, VA with log transformed home range area as response and reproductive season interacting with sex as predictor. Reference categories are sex=female and season=dispersal.

| Covariate | $\beta$ | SE | df | t value | Pr(>\|t\|) |
| --- | --- | --- | --- | --- | --- |
| (Intercept) | 2.370 | 0.202 | 26.95 | 11.755 | < 0.001 |
| Breeding Season | 0.020 | 0.177 | 21.62 | 0.111 | 0.913 |
| Kitten-rearing season | -0.020 | 0.190 | 24.51 | -0.103 | 0.918 |
| Male | 1.098 | 0.236 | 26.69 | 4.658 | < 0.001 |
| Breeding season x male | 0.231 | 0.215 | 22.04 | 1.073 | 0.295 |
| Kitten-rearing season x male | 0.064 | 0.217 | 23.99 | 0.295 | 0.770 |

**Figure 2.**
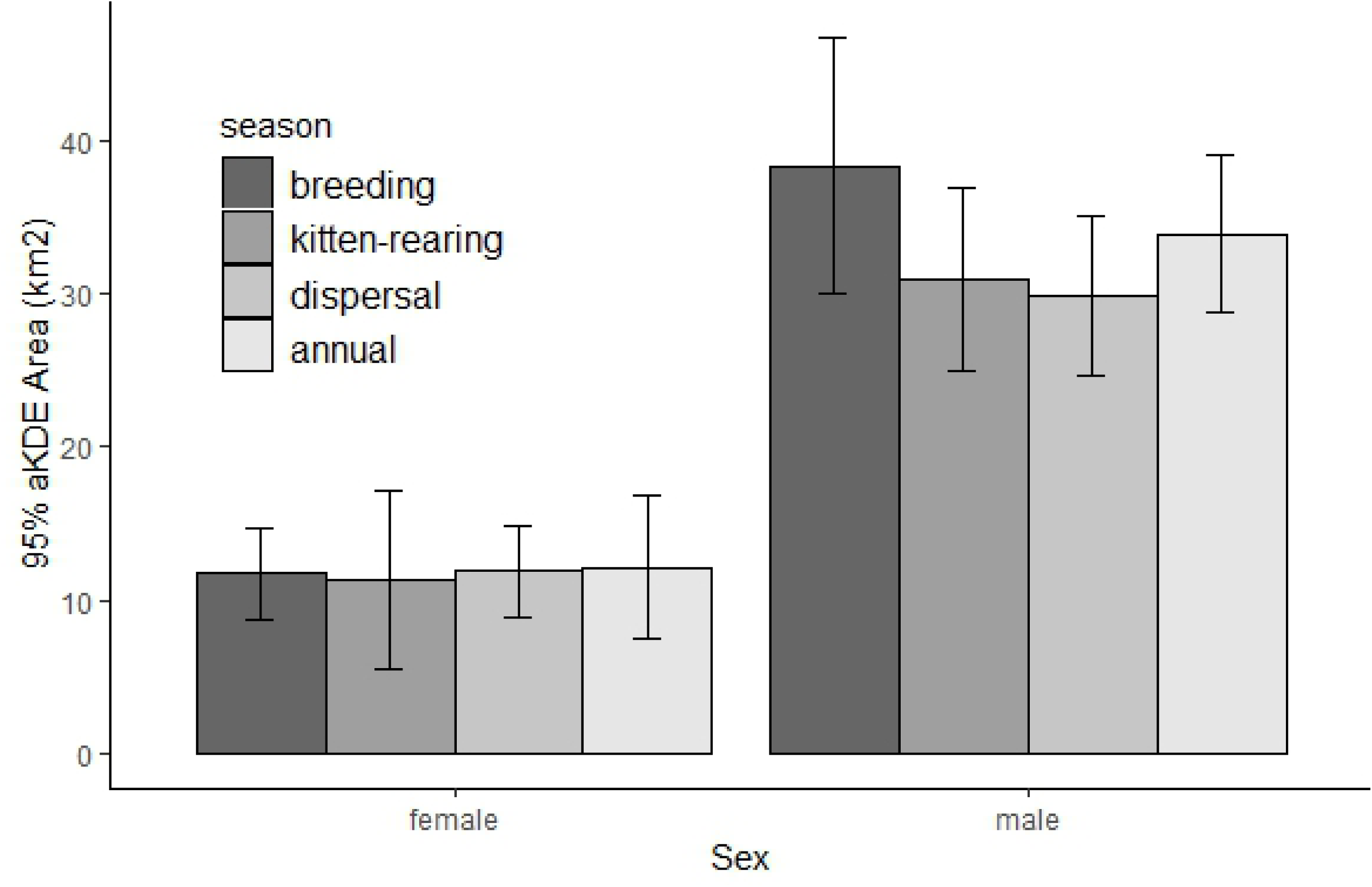
Means and 95% confidence intervals of 95% home ranges of female and male resident bobcats monitored during 2017-2019 in Bath County, VA, for breeding (n=12, 4 females, 8 males), kitten-rearing (n=16, 5 females, 11 males), and dispersal/pre-breeding (n=15, 4 females, 11 males) seasons, and annual (n=16, 5 females, 11 males). Home ranges calculated using the autocorrelated kernel density estimator

We estimated annual movement rates for all bobcats (n=20), and seasonal movement rates for bobcats with at least 1 month of relocations within a given season. Average male movement rates (232.3 ± 12.0 meters/hour) were approximately 1.5 times higher than average female movement rates (154.4 ± 8.9 meters/hour). Male movement rates were higher than female movement rates during all seasons, and female movement rates were significantly higher during the kitten-rearing season while male movement rates were significantly higher during the dispersal season (Table 2, Figure 3).

**Table 2.** Gamma generalized linear mixed-effects model for bobcats monitored during 2017-2019 in Bath County, VA, with movement rates as response and reproductive season interacting with sex as predictor. Reference categories are sex=female and season=kitten-rearing.

| Covariate | $\beta$ | SE | t value | Pr(>\|z\|) |
| --- | --- | --- | --- | --- |
| (Intercept) | 5.127 | 0.099 | 51.867 | < 0.001 |
| Breeding season | -0.150 | 0.072 | -2.093 | 0.036 |
| Dispersal season | -0.149 | 0.057 | -2.605 | 0.009 |
| Male | 0.294 | 0.116 | 2.535 | 0.011 |
| Breeding season x male | 0.082 | 0.078 | 1.060 | 0.289 |
| Dispersal season x male | 0.189 | 0.064 | 2.968 | 0.003 |

**Figure 3.**
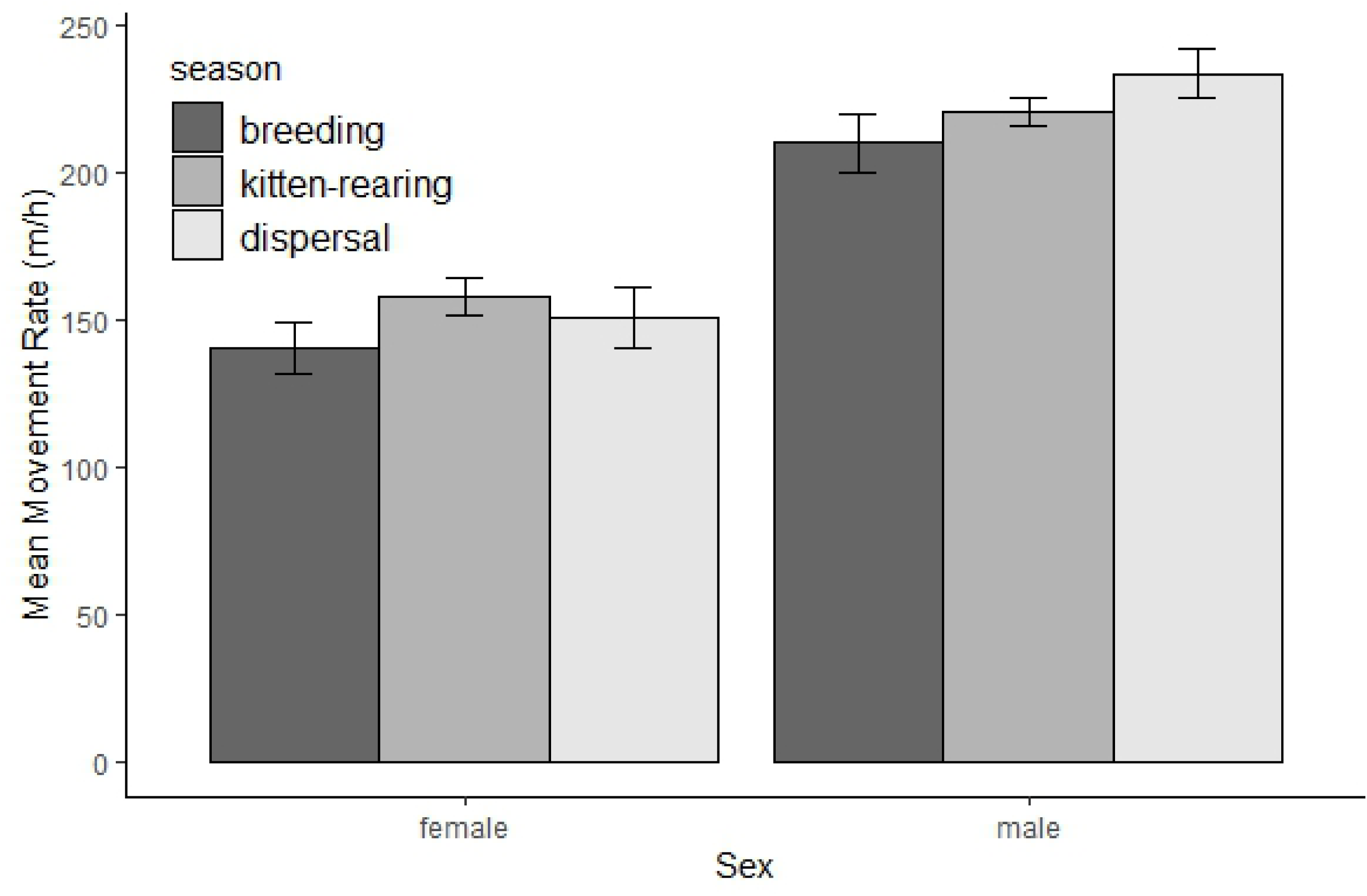
Means and 95% confidence intervals for movement rates of female and male bobcats monitored during 2017-2019 in Bath County, VA, for breeding (n=14, 5 females, 9 males), kitten-rearing (n=17, 4 females, 13 males), and dispersal/pre-breeding (n=15, 4 females, 11 males) seasons. Movement rate is reported as meters moved per hour (m/h).

We conducted 2^nd^ order resource selection for all resident bobcats (12 males, 6 females), which excluded 2 dispersing males. For females distance to deciduous forest, distance to fields, and elevation were the strongest predictors of 2^nd^ order resource selection (Table 3, Figure 4). During all seasons, females selected SAUs that were at higher elevations, closer to deciduous forest, and farther from fields than expected (Table 3, Figure 4). Females exhibited strongest 2^nd^ order selection for deciduous forest during the kitten-rearing season (Table 3, Figure 4). Females exhibited strongest 2^nd^ order avoidance of fields during the kitten-rearing season, weaker avoidance of fields during the dispersal season, and weakest avoidance of fields during the breeding season (Table 3, Figure 4). Females exhibited strongest 2^nd^ order selection for higher elevations during the breeding season, less strong selection for high elevations during the dispersal season, and weakest selection for high elevations during the kitten-rearing season (Table 3, Figure 4). Females exhibited strongest 2^nd^ order selection for mixed forest during the dispersal and breeding seasons, but did not select or avoid mixed forest during the kitten-rearing season (Table 3, Figure 4). Females exhibited 2^nd^ order selection for steeper slopes during the dispersal season, but exhibited 2^nd^ order selection for more gentle slopes during the breeding and kitten-rearing seasons (Table 3, Figure 4).

**Table 3.**
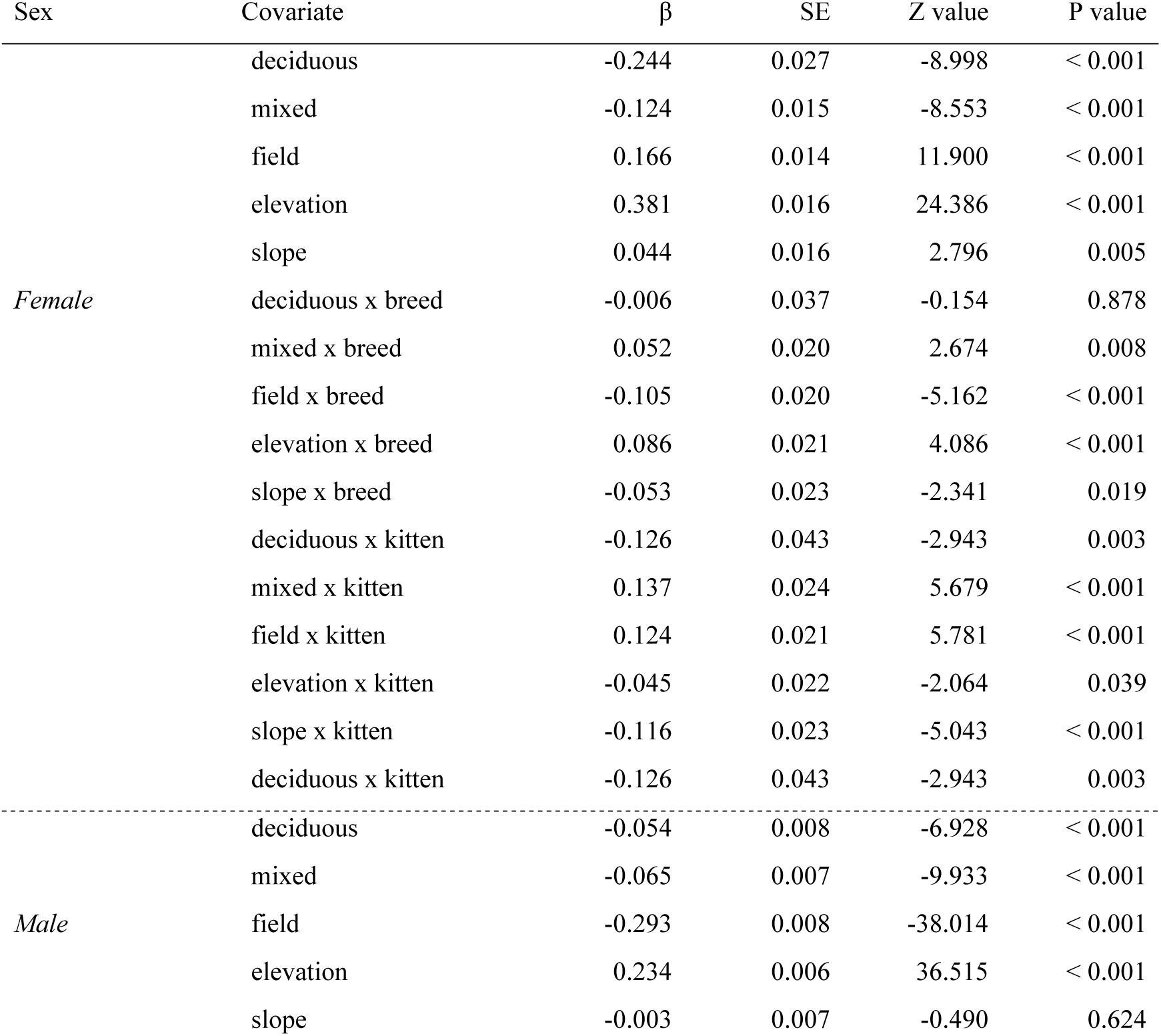
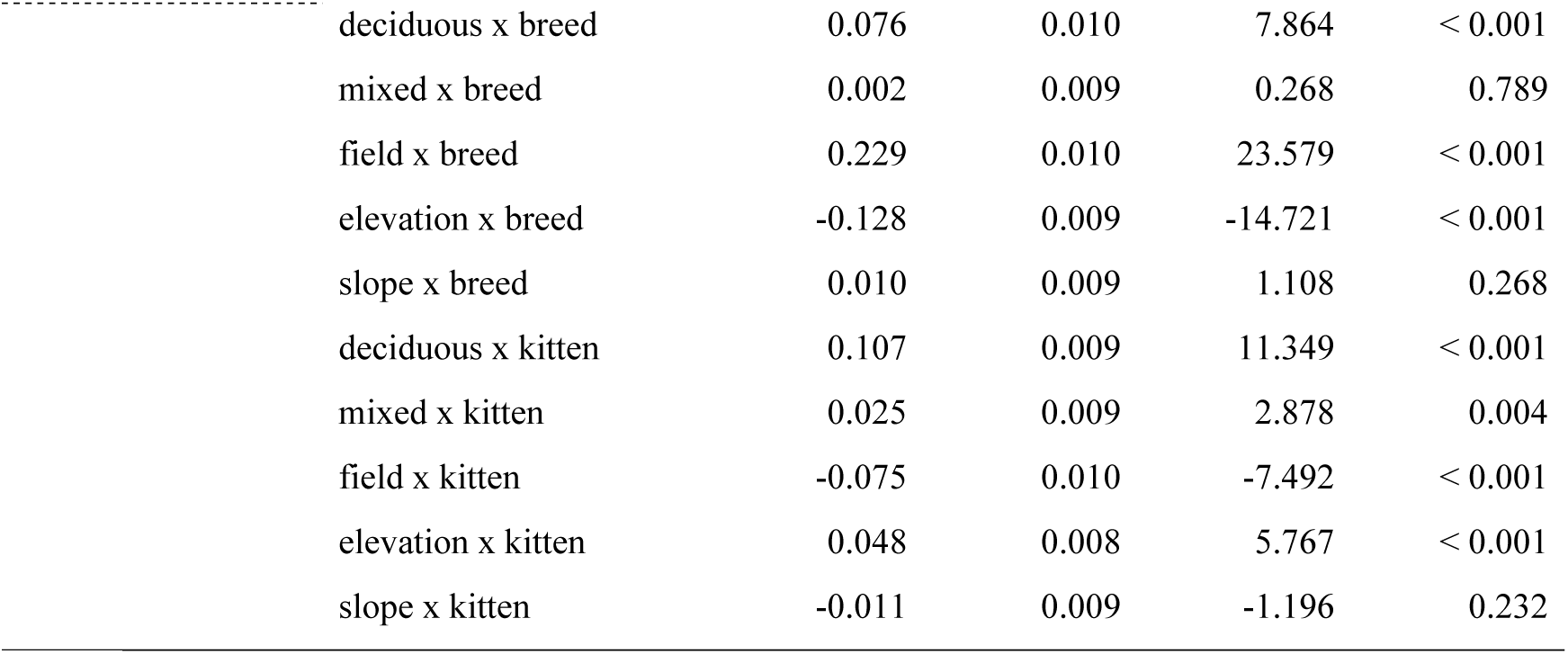
Model results for 2^nd^ order resource selection functions (RSF) for 18 bobcats (12 male, 6 female) collared in Bath County, Virginia in years 2017-2019, including separate models for males and females. RSF models are binomial generalized linear mixed-effects models. Results include β coefficients (β), and standard errors (SE), z values, and p values from Wald tests. Reference category is season = dispersal.

| Sex | Covariate | $\beta$ | SE | Z value | P value |
| --- | --- | --- | --- | --- | --- |
| *Female* | deciduous | -0.244 | 0.027 | -8.998 | < 0.001 |
|  | mixed | -0.124 | 0.015 | -8.553 | < 0.001 |
|  | field | 0.166 | 0.014 | 11.900 | < 0.001 |
|  | elevation | 0.381 | 0.016 | 24.386 | < 0.001 |
|  | slope | 0.044 | 0.016 | 2.796 | 0.005 |
|  | deciduous x breed | -0.006 | 0.037 | -0.154 | 0.878 |
|  | mixed x breed | 0.052 | 0.020 | 2.674 | 0.008 |
|  | field x breed | -0.105 | 0.020 | -5.162 | < 0.001 |
|  | elevation x breed | 0.086 | 0.021 | 4.086 | < 0.001 |
|  | slope x breed | -0.053 | 0.023 | -2.341 | 0.019 |
|  | deciduous x kitten | -0.126 | 0.043 | -2.943 | 0.003 |
|  | mixed x kitten | 0.137 | 0.024 | 5.679 | < 0.001 |
|  | field x kitten | 0.124 | 0.021 | 5.781 | < 0.001 |
|  | elevation x kitten | -0.045 | 0.022 | -2.064 | 0.039 |
|  | slope x kitten | -0.116 | 0.023 | -5.043 | < 0.001 |
|  | deciduous x kitten | -0.126 | 0.043 | -2.943 | 0.003 |
| *Male* | deciduous | -0.054 | 0.008 | -6.928 | < 0.001 |
|  | mixed | -0.065 | 0.007 | -9.933 | < 0.001 |
|  | field | -0.293 | 0.008 | -38.014 | < 0.001 |
|  | elevation | 0.234 | 0.006 | 36.515 | < 0.001 |
|  | slope | -0.003 | 0.007 | -0.490 | 0.624 |

|  |  |  |  |  |
| --- | --- | --- | --- | --- |
| deciduous x breed | 0.076 | 0.010 | 7.864 | < 0.001 |
| mixed x breed | 0.002 | 0.009 | 0.268 | 0.789 |
| field x breed | 0.229 | 0.010 | 23.579 | < 0.001 |
| elevation x breed | -0.128 | 0.009 | -14.721 | < 0.001 |
| slope x breed | 0.010 | 0.009 | 1.108 | 0.268 |
| deciduous x kitten | 0.107 | 0.009 | 11.349 | < 0.001 |
| mixed x kitten | 0.025 | 0.009 | 2.878 | 0.004 |
| field x kitten | -0.075 | 0.010 | -7.492 | < 0.001 |
| elevation x kitten | 0.048 | 0.008 | 5.767 | < 0.001 |
| slope x kitten | -0.011 | 0.009 | -1.196 | 0.232 |

**Figure 4.**
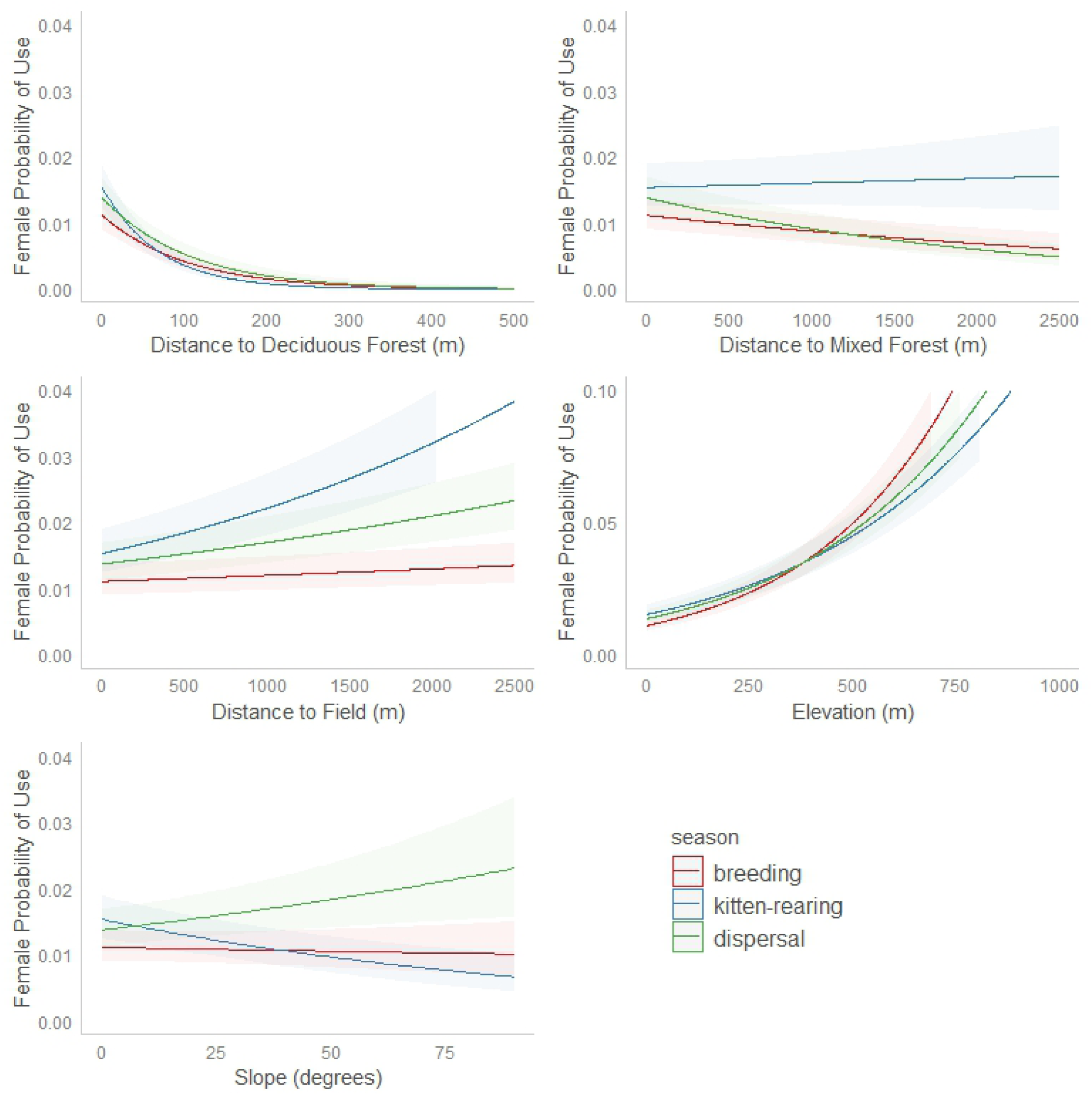
Relative probability of 2^nd^ order selection with 95% confidence intervals for female bobcats monitored during 2017-2019 in Bath County, VA, for breeding (n=5), kitten-rearing (n=4), and dispersal/pre-breeding (n=4) seasons.

For males, distance to fields and elevation were the strongest predictors of 2^nd^ order resource selection (Table 3, Figure 4). During all seasons, males selected SAUs that were closer to fields and at higher elevations than expected (Table 3, Figure 5). Males exhibited strongest 2^nd^ order selection for fields during the kitten-rearing season and weakest selection for fields during the breeding season (Table 3, Figure 5). Males exhibited weakest 2^nd^ order selection for high elevations during the breeding season compared to dispersal and kitten-rearing seasons (Table 3, Figure 5). Males exhibited 2^nd^ order selection for mixed forest during all seasons, but this selection was weakest during the kitten-rearing season, following a similar pattern as females (Table 3, Figure 5). Males exhibited 2^nd^ order selection for deciduous forest during the dispersal season, but avoided deciduous forest during breeding and kitten-rearing seasons (Table 3, Figure 5). Slope was not a significant predictor of male 2^nd^ order resource selection (Table 3, Figure 5).

**Figure 5.**
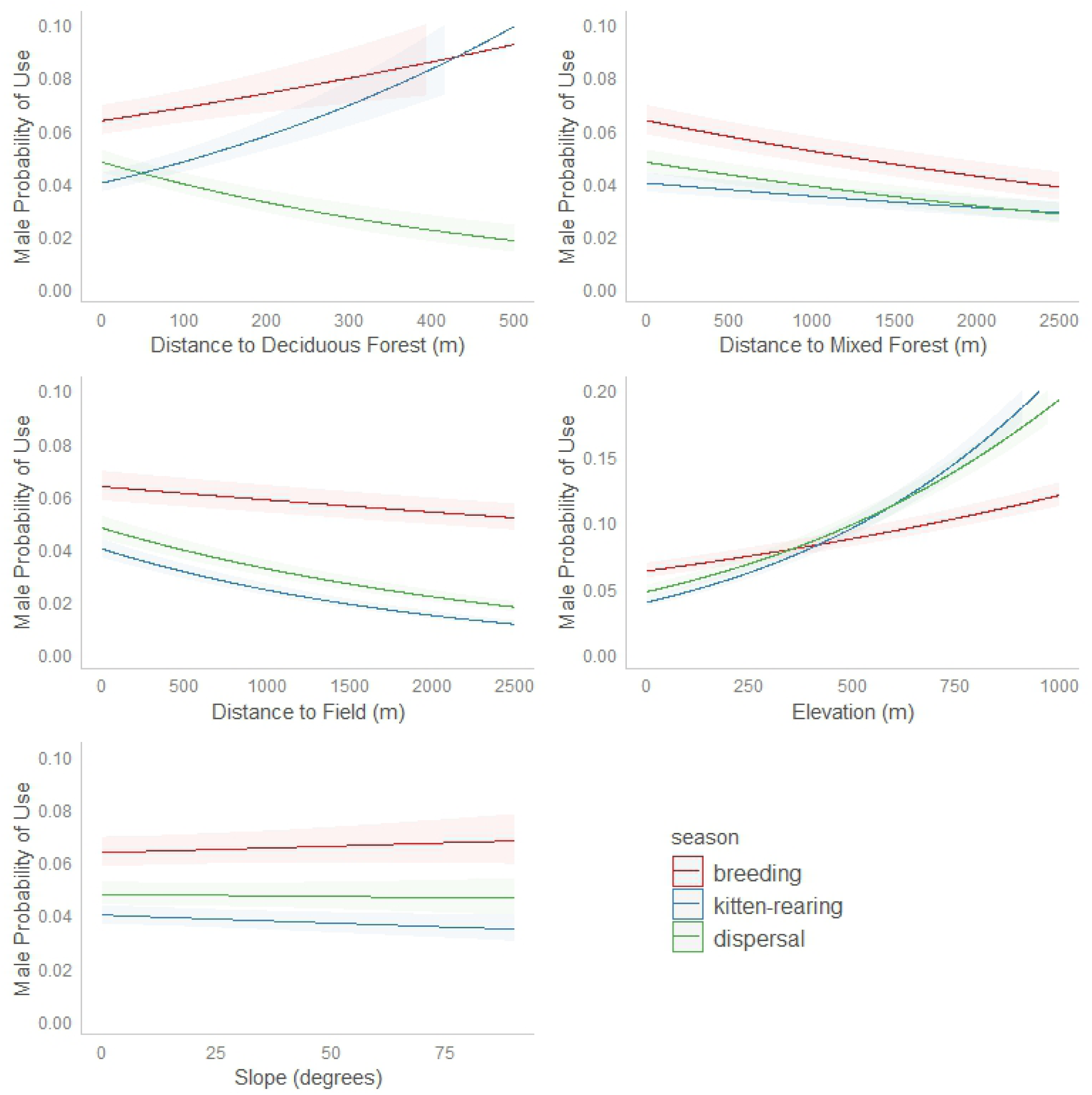
Relative probability of 2^nd^ order selection with 95% confidence intervals for male bobcats monitored during 2017-2019 in Bath County, VA, for breeding (n=9), kitten-rearing (n=13), and dispersal/pre-breeding (n=11) seasons.

We examined 3^rd^ order resource selection across seasons for male bobcats with fix success of ≥85% (n=7). Distance to deciduous forest, distance to fields, elevation, and slope were all significant predictors of male 3^rd^ order resource selection (Table 4, Figure 6). Male bobcats exhibited 3^rd^ order selection for deciduous forest during all seasons (Table 4, Figure 6). Male bobcats exhibited 3^rd^ order selection for fields during all seasons, but this selection was stronger during the kitten-rearing season (Table 4, Figure 6). Male bobcats exhibited 3^rd^ order selection for high elevations during all seasons, with the strongest selection for high elevations during the dispersal season and weakest selection during the breeding season (Table 4, Figure 6). Lastly, male bobcats exhibited 3^rd^ order selection for gentle slopes during the kitten-rearing and dispersal seasons, but selected for steeper slopes during the breeding season (Table 4, Figure 6).

**Table 4.**
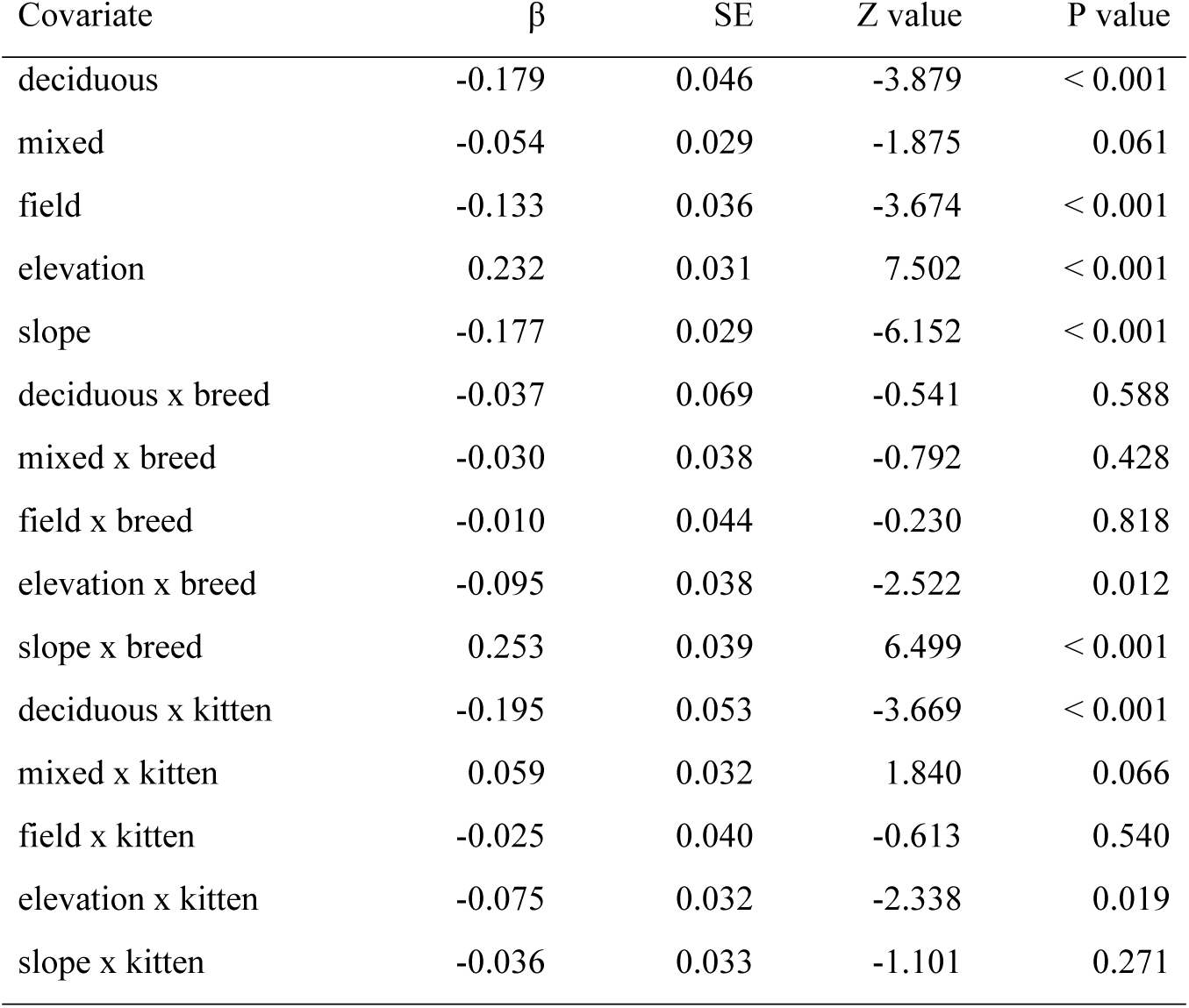
Model results for 3^rd^ order resource selection functions (RSF) for male bobcats (n=7) collared in Bath County, Virginia in 2018-2019. RSF model is a binomial generalized linear mixed-effects model. Results include β coefficients (β), and standard errors (SE), z values, and p values from Wald tests. Reference category is season = dispersal.

| Covariate | $\beta$ | SE | Z value | P value |
| --- | --- | --- | --- | --- |
| deciduous | -0.179 | 0.046 | -3.879 | < 0.001 |
| mixed | -0.054 | 0.029 | -1.875 | 0.061 |
| field | -0.133 | 0.036 | -3.674 | < 0.001 |
| elevation | 0.232 | 0.031 | 7.502 | < 0.001 |
| slope | -0.177 | 0.029 | -6.152 | < 0.001 |
| deciduous x breed | -0.037 | 0.069 | -0.541 | 0.588 |
| mixed x breed | -0.030 | 0.038 | -0.792 | 0.428 |
| field x breed | -0.010 | 0.044 | -0.230 | 0.818 |
| elevation x breed | -0.095 | 0.038 | -2.522 | 0.012 |
| slope x breed | 0.253 | 0.039 | 6.499 | < 0.001 |
| deciduous x kitten | -0.195 | 0.053 | -3.669 | < 0.001 |
| mixed x kitten | 0.059 | 0.032 | 1.840 | 0.066 |
| field x kitten | -0.025 | 0.040 | -0.613 | 0.540 |
| elevation x kitten | -0.075 | 0.032 | -2.338 | 0.019 |
| slope x kitten | -0.036 | 0.033 | -1.101 | 0.271 |

**Figure 6.**
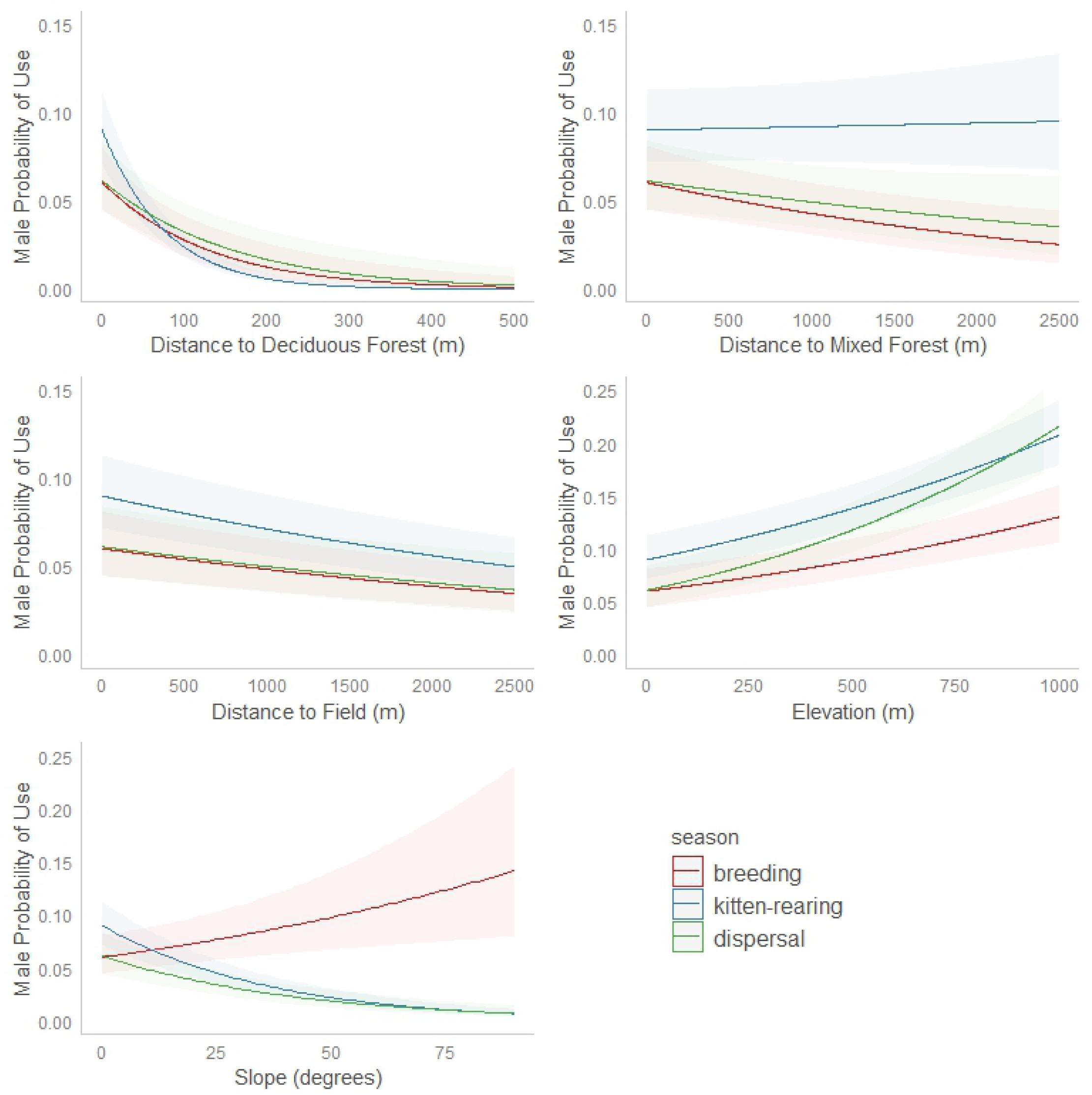
Relative probability of 3^rd^ order selection with 95% confidence intervals for male bobcats monitored during 2017-2019 in Bath County, VA, for breeding (n=5), kitten-rearing (n=7), and dispersal/pre-breeding (n=7) seasons.

## DISCUSSION

Bobcat space use is primarily driven by the need to reproduce and acquire prey. Thus, differences in reproductive pressures between sexes likely explain most of the variation between sexes that we observed, and reproductive seasons and seasonal fluctuations in prey availability likely explain most of the variation among seasons.

Despite the fact that the central Appalachian Mountains experience climatic conditions similar to more northern climates due to high elevations, our bobcat home range estimates are more similar in size to estimates from southern study areas, and we did not observe seasonal changes in home range size as is often found in northern latitudes. Bobcat home range estimates from northern New York and Maine were 3-4 times larger than we observed [21,46], possibly because our study area does not experience severe winters or sustained, deep snow as found farther north. Bobcats are poorly adapted to foraging in deep snow [28,47]. In areas with severe winters and sustained snow, bobcat home ranges can increase drastically during winter months [48,49]

Male bobcats in our study had larger home ranges than females, in accordance with nearly all bobcat space use studies [13]. This supports the prediction that males seek to maximize breeding opportunities by extending their space use to overlap multiple female home ranges, whereas females use space more strictly to acquire resources [12]. It is unlikely that metabolic demands explain this sex difference, as the disparity in home range size between sexes (males approximately 3 times larger) far exceeded the disparity in body size (males approximately 1.5 times larger). Interestingly, SAU size for males did not differ between breeding and non-breeding seasons, indicating that pressure to maximize reproductive success drives male space use even at times distant from active breeding. Perhaps the costs of reestablishing large territories immediately prior to the breeding season are greater than maintaining them throughout the year. Likewise, female SAU size did not vary across seasons, even though metabolic demands increase during kitten-rearing. Instead it appears that bobcats in our study area use home ranges more intensively during certain periods, indicated by seasonal increases in movement rates for both sexes.

The increased female movement rates we observed during the kitten-rearing season are likely due to the need to increase foraging but also attend to young, resulting in frequent movements between den sites and foraging sites. Female bobcat activity rates have been found to increase during kitten rearing [50]. The increased male movement rates we observed during dispersal season may be due to transient bobcats seeking home ranges prior to the breeding season, and increased territorial marking and patrolling by resident males in response to transients in proximity. Physical territorial conflicts between bobcats are thought to be largely avoided by communicating through urine spraying, feces deposition, physical scrapes, olfactory investigation, and vocalizations [41]. Male movement rates were greater than female movement rates throughout the year, which may be related to their overall larger home range size. With larger home ranges, males must move greater distances to patrol territories and move between multiple female home ranges.

Both male and female bobcats strongly selected home ranges at higher elevations. In the topography of the Valley and Ridge province, this is a telling trend, as ridges are largely forested while most development and agriculture occur in valley bottoms. However, other resource selection patterns differed between sexes. Male bobcats exhibited 2^nd^ order selection for fields and avoidance of deciduous forest, whereas female bobcats contrastingly exhibited 2^nd^ order avoidance of fields and selection of deciduous forest. These opposing trends may be explained by the valley and ridge topography of the study area, combined with differing reproductive pressures. Female home ranges almost exclusively occurred within a single ridge, whereas male home ranges often contained multiple ridges (Figure 1), perhaps reflecting attempts to overlap multiple female home ranges. In moving between ridges, male bobcats must cross valleys and therefore encompass fields within their home ranges. Chamberlain et al. [15] found male bobcat resource selection to vary by scale, and suggested that males may select home ranges primarily based on the spatial distribution of females, which aligns with the space use trends we observed.

At the 3^rd^ order, male bobcats strongly selected for deciduous forest and high elevations during all seasons, indicating selection of forested ridges within home ranges. Male bobcats also selected for fields within home ranges. Bobcats are known to select for both forest habitat and dense understory vegetation [21,24,25,26]. In the closed-canopy forests found in the Appalachian Mountains, much of the dense understory is found along forest-field edges. Considering our findings of male resource selection across scales, it appears that 2^nd^ order selection of male bobcats was influenced more by the spatial distribution of females, whereas 3^rd^ order selection was influenced more by resource acquisition.

As bobcat prey selection shifts throughout the year, habitat selection may shift accordingly [15,29]. The availability of prey and cover can change drastically across seasons in our system, which may explain the seasonal resource selection shifts that we observed. In our study area, the season we have termed kitten-rearing also overlaps with the growing season, and the seasons we have termed dispersal and breeding overlap with the dormant season. During the growing season, herbaceous plants and deciduous trees provide much of the cover across the landscape. White-tailed deer fawns, juvenile cottontail rabbits, and young of many other common bobcat prey species are abundant and relatively easily acquired during this time. In contrast, cover provided by herbaceous plants and deciduous foliage is lacking during the dormant season, and prey abundance is generally lower due to the lack of newborn prey.

Both sexes exhibited 2^nd^ order selection for mixed forest during the dispersal and breeding seasons, but they either exhibited weaker selection for, or did not select for, mixed forest during the kitten-rearing season. Other studies have found bobcat selection for conifers during winter months, which has been attributed to relatively higher prey availability during winter [29,51,52]. We found male bobcats increasingly selected for fields during the kitten-rearing season, which overlaps with the growing season. Female white-tailed deer have been found to select parturition sites and bed sites in areas with high visual obscurity, which often occurs along forest-field edges (Shuman et al. 2018). Likewise, cottontail rabbits select for grassy areas and dense vegetation found in overgrown fields and along field edges [54,55]. Male bobcats also exhibited 2^nd^ order selection for deciduous forest only during the dispersal season, which overlaps with the peak of hard mast production. Squirrels are the most common diet item of bobcats in this study area [8], and both gray squirrels (*Sciurus carilonensis)* and fox squirrels (*Sciurus niger)* exhibit peaks in foraging behavior during this time [56,57,58].

Our findings build on the scarce knowledge of bobcat spatial ecology in the central Appalachian Mountains, particularly the absence of published bobcat resource selection data for the region. Further, this is the first study to utilize GPS collars on bobcats in the central Appalachian Mountains. This study has illuminated difficulties in conducting GPS telemetry in the dense canopy and rugged landscape of Appalachia, particularly on bobcats due to their size and life history. The main limitation of this study was the poor fix success of some GPS collars, and the compounding effects with the already small sample size that prohibited 3^rd^ order resource selection analysis for females. Despite this fact, GPS relocations certainly exceeded VHF-telemetry samples from previous nearby studies in sample size, frequency, and locational accuracy; allowing a more robust examination of space use and resource selection.

### Management and Research Implications

During the early 20^th^ century, bobcat populations persisted in the mountainous western portion of Virginia, while populations in most eastern portions of the state were extirpated or heavily reduced (VDGIF unpublished data). Thus, the Appalachian Mountains acted as refugia during the period of extirpation, likely due to their rugged, inaccessible terrain. Our findings that bobcats select for higher elevations and deciduous forest (i.e. ridges) in the systematic patchwork of the Valley and Ridge province indicate the importance of these features today. Conversion of valley bottoms from rich riparian forests to agriculture and development has probably shifted bobcat space use from historical patterns, leading to increased use of ridges. Due to their rugged terrain, most ridges are forested, public land. The long, undeveloped ridges of the Valley and Ridge province facilitate resident bobcat movement and potentially act as dispersal corridors, particularly in areas with more intensely-developed valley bottoms. Bobcat selection for deciduous forest that we observed, even though deciduous forest is highly available (roughly 90% of the study area), highlights the importance of forest cover. When considering the potential effects of future land use in the region, particularly the development of ridges, managers should consider the importance of landscape features to bobcats. Although forested habitat is an important component of bobcat habitat, dense understory is also an important consideration. Using timber harvest and prescribed fire, managers can increase the availability of dense understory cover in those forests (McNitt et al. under review).

Male bobcats exhibited selection for fields, and this selection was strongest during the kitten-rearing season at both scales. The kitten-rearing season overlaps with the time period in which juvenile prey are present on the landscape, and it is possible that prey species select areas of high visual obscurity near field edges to birth and raise young, as is found in white-tailed deer [53,59]. Thus, selection of fields during summer months may indicate that males increase foraging along field edges at this time. Further research should investigate where bobcat predation is most likely to occur, and if field edges are areas of high predation risk.

Future research should consider the variation in bobcat space use across sex and season. For example, male bobcats will be more detectable due to larger home ranges and greater movement rates. Particularly noninvasive research, such as population and diet sampling, should account for these differences in detectability when considering study design, data analysis, and interpretation of results. Research into bobcat predation, and how prey selection may vary between sexes or across seasons, may further explain the seasonal shifts in resource selection that we observed and potential implications to prey species.

## Acknowledgements

We appreciated extensive logistical support from the Virginia Department of Game and Inland Fisheries, US Forest Service, and The Nature Conservancy. Thanks to N. Simmons of The Nature Conservancy, and S. Tanguay and E. McNichols of the USFS. We appreciated assistance with analysis from D. Crawford. We thank D. Morin, W.H. Ellsworth, J. Zanon, B. George, C. Satter, C. Rowe, A. Hilborn, and C. Osorio for assistance in the field. Thanks to many members of the public in Bath County, VA for support in the field.

## References

1. Manly BFJ, McDonald LL, Thomas DL, McDonald TL, Erickson W. Resource selection by animals: statistical design and analysis for field studies. Kluwer Academic Publishers. 2002.

2. Millspaugh, JJ, Marzluff JM. Radio tracking and animal populations. San Diego, Academic Press. 2001.

3. Börger L, Dalziel BD, Fryxell JM. Are there general mechanisms of animal home range behaviour? A review and prospects for future research: home range modelling. Ecology Letters. 2008;11(6):637–650.

4. Boyce MS, Vernier PR, Nielsen SE, Schmiegelow FKA. Evaluating resource selection functions Ecological Modelling. 2002;157(2-3):281–300

5. Johnson DH. The comparison of usage and availability measurements for evaluating resource preference. Ecology. 1980;61(1):65–71.

6. Roberts NM, Crimmins SM. Bobcat Population Status and Management in North America: Evidence of Large-Scale Population Increase. Journal of Fish and Wildlife Management. 2010;1(2):169–174. doi:10.3996/122009-JFWM-026

7. Conner, ML, Leopold, BD, and MJ Chamberlain. Multivariate habitat models for bobcats in southern forested landscapes In: Woolf, A, Nielsen, CK, Bluett, RD, (Eds), Proceedings of a symposium on Current Bobcat Research and Implications for Management. pp 51–55. 2001.

8. Morin DJ, Higdon SD, Holub JL, Montague DM, Fies ML, Waits LP, Kelly MJ. Bias in carnivore diet analysis resulting from misclassification of predator scats based on field identification: bias and uncertainty in carnivore scat identification. Wildlife Society Bulletin. 2016;40(4):669–677.

9. Morin DJ, Kelly MJ, Waits LP. Monitoring coyote population dynamics with fecal DNA and spatial capture-recapture: Estimating Coyote Population Parameters. The Journal of Wildlife Management. 2016;80(5):824–836. doi:10.1002/jwmg.21080

10. Morin DJ, Waits LP, McNitt DC, Kelly MJ. Efficient single-survey estimation of carnivore density using fecal DNA and spatial capture-recapture: a bobcat case study. Population Ecology. 2018;60(3):197–209. doi:10.1007/s10144-018-0606-9

11. Anderson EM, Lovallo MJ. Bobcat and lynx Wild mammals of North America: biology, management, and economics. Baltimore: Johns Hopkins Press; 2003. pp. 758–786.

12. Sandell M. The mating tactics and spacing patterns of solitary carnivores. In Carnivore Behavior, Ecology, and Evolution (Ed by Gittleman JL). Ithaca, Cornell University Press. 1989; pp 164–182.

13. Ferguson AW, Currit NA, Weckerly FW. Isometric scaling in home-range size of male and female bobcats (Lynx rufus). Canadian Journal of Zoology 2009;87(11): 1052–1060.

14. Bailey TN. Social organization in a bobcat population. Journal of Wildlife Management. 1974:38(3): 435–446

15. Chamberlain MJ, Leopold BD, Conner LM. Space use, movements and habitat selection of adult bobcats (Lynx rufus) in central Mississippi. American Midland Naturalist. 2003;149(2):395–405.

16. Kitchings JT, Story JD. Movements and dispersal of bobcats in east Tennessee. Journal of Wildlife Management. 1984;48(3):957–961.

17. Knowles PR. Home range size and habitat selection of bobcats, Lynx rufus, in north-central Montana. Canadian Field-Naturalist. 1985;99: 6–12.

18. Abouelezz HG, Donovan TM, Mickey RM, Murdoch JD, Freeman M, Royar K. Landscape composition mediates movement and habitat selection in bobcats (Lynx rufus): implications for conservation planning. Landscape Ecology. 2018;33: 1301–1318.

19. Elizalde-Arellano CJ, López-Vidal C, Hernández L, Laundré JW, Cervantes FA, Alonso-Spilsbury M. Home range size and activity patterns of bobcats (Lynx rufus) in the southern part of their range in the Chihuahuan Desert, Mexico. The American Midland Naturalist. 2012;168(2): 247–264

20. Rockhill AP, DePerno CS, Powell RA. The Effect of Illumination and Time of Day on Movements of Bobcats (Lynx rufus). PLoS ONE. 2013;8(7):e69213. doi:10.1371/journal.pone.0069213

21. Litvaitis JA, Sherburne JA, Bissonette JA. Bobcat habitat use and home range size in relation to prey density. Journal of Wildlife Management. 1986;50(1): 110–117.

22. Knick ST. Ecology of bobcats relative to exploitation and a prey decline in southeastern Idaho. Wildlife Monographs. 1990. pp 3–42.

23. Godbois IA, Conner LM, Warren RJ. Space-use patterns of bobcats relative to supplemental feeding of northern bobwhites. Journal of Wildlife Management. 2004;68(3): 514–518.

24. Kolowski JM, Woolf A. Microhabitat use by bobcats in southern Illinois. Journal of Wildlife Management. 2002;66(3):822.

25. Mosby CE, Grovenburg TW, Klaver RW, Schroeder GM, Schmitz LE, and Jenks JA. Microhabitat selection by bobcats in the Badlands and Black Hills of South Dakota, USA: A comparison of prairie and forested habitats. The Prairie Naturalist. 2012;44(1): 47–57.

26. Tucker SA, Clark WR, Gosselink TE. Space Use and Habitat Selection by Bobcats in the Fragmented Landscape of South-Central Iowa. Journal of Wildlife Management. 2008;72(5):1114–1124. doi:10.2193/2007-291

27. Cochrane JC, Kirby JD, Jones IG, Conner ML, Warren RJ. Spatial Organization of adult bobcats in a longleaf pine-wiregrass ecosystem in Southwestern Georgia. Southeastern Naturalist. 2006;5(4): 711–724.

28. Koehler GM, Hornocker MG. Influences of seasons on bobcats in Idaho. Journal of Wildlife Management. 1989;53(1): 197–202.

29. Lovallo MJ, Anderson EM. Bobcat (Lynx rufus) home range size and habitat use in Northwest Wisconsin. American Midland Naturalist. 1996;135(2):241. doi:10.2307/2426706

30. Rucker RA, Kennedy ML, Heidt GA, Harvey MJ. Population density, movements, and habitat use of bobcats in Arkansas. Southwestern Naturalist. 1989;34(1): 101–108.

31. Lancia RA. Summer movement patterns and habitat use by bobcats on Croatan National Forest, North Carolina. Cats of the World: Biology, Conservation, and Management. 1986. pp 425–436.

32. Kitchings JT, Story JD. Movements and dispersal of bobcats in east Tennessee. Journal of Wildlife Management. 1984;48(3):957–961.

33. Whitaker J, Fredrick RB, Edwards LT. Home-range size and overlap of eastern Kentucky bobcats. In Proceedings of the Southeastern Association of Fish and Wildlife Agencies. 1987;41: 417–423

34. Progulske DR. Game Animals Utilized as Food by the Bobcat in the Southern Appalachians. The Journal of Wildlife Management. 1955;19(2):249. doi:10.2307/3796859

35. Jackson DL, Gluesing EA, Jacobson HA. Dental eruption in bobcats. Journal of Wildlife Management. 1988;52(3): 515–517.

36. Stys ED, Leopold BD. Reproductive biology and kitten growth of captive bobcats in Mississippi. Proceedings of the annual conference Southeastern Association of Fish and Wildlife Agencies. 1993;47: 80–89.

37. Winegarner CE, Winegarner MS. Reproductive history of a bobcat. Journal of Mammalogy. 1982;63(4):680–681. doi:10.2307/1380281

38. Fleming CH, Fagan WF, Mueller T, Olson KA, Leimgruber P, Calabrese JM. 2015 Rigorous home range estimation with movement data: a new autocorrelated kernel density estimator. Ecology 96(5):1182–1188.

39. Fleming CH, Calabrese JM. ctmm: Continuous-Time Movement Modeling R package version 052. 2018.

40. Fleming CH, Sheldon D, Fagan WF, Leimgruber P, Mueller T, Nandintsetseg D … et al. Correcting for missing and irregular data in home-range estimation. Ecological Applications. 2018;28(4):1003–1010.

41. Allen, M. L., C. F. Wallace, and C. C. Wilmers. Patterns in bobcat (Lynx rufus) scent marking and communication behaviors. Journal of Ethology. 2015;33: 9–14.

42. Bates, D, M Mächler, B Bolker, and S Walker. Fitting linear mixed-effects models using lme4 Journal of Statistical Software. 2015. doi:10.18637/jss.v067.i01

43. Kuznetsova A, Brockhoff PB, Christensen RHB. “lmerTest Package: Tests in Linear Mixed Effects Models”. Journal of Statistical Software 2017;82(13): 1–26.

44. Katnik DD, Wielgus RB. Landscape proportions versus Monte Carlo simulated home ranges for estimating habitat availability. Journal of Wildlife Management. 2005;69(1): 20–32.

45. Gillies CS, Hebblewhite M, Nielsen SE, Krawchuk MA, Aldridge CL, Frair JL … et al. Application of random effects to the study of resource selection by animals: random effects in resource selection. Journal of Animal Ecology. 2006;75: 887–898.

46. Fox LB. Ecology and population biology of the bobcat (*Felis rufus*) in New York. PhD Dissertation, State University of New York College of Environmental Science and Forestry. 1990.

47. Gooliaff T, Hodges KE. Historical distributions of bobcats (*Lynx rufus*) and Canada lynx (*Lynx canadensis*) suggest no range shifts in British Columbia, Canada. Canadian Journal of Zoology. 2018;96(12):1299–1308.

48. Donovan TM, Freeman M, Abouelezz H, Royar K, Howard A, and Mickey R. Quantifying home range habitat requirements for bobcats (Lynx rufus) in Vermont, USA. Biological Conservation. 2011;144(12): 2799–2809.

49. Reed GC, Litvaitis JA, Ellingwood M, et al. Describing habitat suitability of bobcats (Lynx rufus) using several sources of information obtained at multiple spatial scales. Mammalian Biology. 2017;82:17–26. doi:10.1016/j.mambio.2016.10.002

50. Chamberlain MJ. Dietary patterns of sympatric bobcats and coyotes in central Mississippi. In Proceedings of the Annual Conference of the Southeastern Association of Fish and Wildlife Agencies. 1999;53: 204–219.

51. McCord CM. Selection of winter habitat by bobcats (Lynx rufus) on the Quabbin Reservation, Massachusetts. Journal of Mammalogy. 1974;55(2):428–437.

52. Rollings CT. Habits, foods and parasites of the bobcat in Minnesota. Journal of Wildlife Management. 1945; 9:131–145.

53. Shuman RM, Cherry MJ, Dutoit EA, Simoneaux TN, Miller KV, Chamberlain MJ. Resource Selection by Parturient and Post-parturient White-tailed Deer and their Fawns. Journal of the Southeastern Association of Fish and Wildlife Agencies 2018;5: 78–84.

54. Althoff D P, Storm GL, Dewalle D.R.. Daytime habitat selection by cottontails in central Pennsylvania. The Journal of Wildlife Management. 1997;61(2): 450.

55. Bond BT, Burger Jr. LW, Leopold BD, Jones JC, Godwin KD. Habitat use by cottontail rabbits across multiple spatial scales in Mississippi. The Journal of wildlife management. 2002;66(4): 1171–1178.

56. Fox JF. Adaptation of gray squirrel behavior to autumn germination by white oak acorns Evolution. 1982;36(4): 800–809.

57. Short HL, Duke WB. Seasonal food consumption and body weights of captive tree squirrels. Journal of Wildlife Management.. 1971;35(3): 435–439.

58. Spritzer MD. Diet, microhabitat use and seasonal activity patterns of gray squirrels (*Sciurus carolinensis*) in hammock and upland pine forest. American Midland Naturalist. 2002;148(2): 271–282.

59. Cherry MJ, Warren RJ, Conner LM. Fire-mediated foraging tradeoffs in white-tailed deer. Ecosphere. 2017;8(4):e01784. doi:10.1002/ecs2.1784

